# Direct genome sequencing of respiratory viruses from low viral load clinical specimens using target capture sequencing technology

**DOI:** 10.1101/2024.04.05.588295

**Authors:** Nobuhiro Takemae, Yumani Kuba, Kunihiko Oba, Tsutomu Kageyama

## Abstract

The use of metagenomic next-generation sequencing technology to obtain complete viral genome sequences directly from clinical samples with low viral load remains challenging—especially in the case of respiratory viruses—due to the low copy number of viral versus host genomes. To overcome this limitation, target capture sequencing for the enrichment of specific genomes has been developed and applied for direct genome sequencing of viruses. However, as the efficiency of enrichment varies depending on the probes, the type of clinical sample, etc., validation is essential before target capture sequencing can be applied to clinical diagnostics. Here we evaluated the utility of target capture sequencing with a comprehensive viral probe panel for clinical respiratory specimens collected from patients diagnosed with SARS-CoV-2 or influenza type A. We focused on clinical specimens containing low copy numbers of viral genomes. Target capture sequencing yielded approximately 180- and 2000-fold higher read counts of SARS-CoV-2 and influenza A virus, respectively, than metagenomic sequencing when the RNA extracted from specimens contained 59.3 copies/µL of SARS-CoV-2 or 544 copies/µL of influenza A virus, respectively. In addition, the target capture sequencing identified sequence reads in all SARS-CoV-2- or influenza type A-positive specimens with <26 RNA copies/µL, some of which also yielded >70% of the full-length genomes of SARS-CoV-2 or influenza A virus. Furthermore, the target capture sequencing using comprehensive probes identified co-infections with viruses other than SARS-CoV-2, suggesting that this approach will not only detect a wide range of viruses, but also contribute to epidemiological studies.

## Introduction

Next-generation sequencing (NGS) techniques have proven to be an indispensable tool for monitoring and controlling emerging pathogens, as demonstrated by the SARS-CoV-2 pandemic (1). In addition, high-throughput and parallel nucleotide sequence analysis by NGS is enabling metagenomic sequencing, which could provide comprehensive genome sequences present in specimens (2). However, metagenomic sequencing has a lower detection sensitivity for viral genomes than traditional diagnostic methods such as PCR because the copy number of viral genomes is much lower than that of host and bacterial genomes in clinical specimens (3). Therefore, obtaining full viral genome sequences directly from clinical specimens with low viral load is still challenging, especially for respiratory viruses, and full viral genome sequencing is often performed after virus isolation.

To overcome the weaknesses of NGS, an enrichment of certain genomes using amplicon-based sequencing (or the tiling amplicon method) or target capture sequencing (or hybridization capture sequencing) has been developed and applied for direct genome sequencing of viruses in clinical or environmental samples (3–5). In the amplicon-based sequencing, single or multiple regions of the target genomes can be amplified using from one set up to many sets of specific primers, providing a highly sensitive and economical method (4). This approach has therefore been used to obtain whole genome sequences of SARS-CoV-2 from clinical specimens worldwide (6–8). However, because whole viral genome sequences are no longer available if unexpected mutations occur in the primer regions, the primers must be updated frequently, especially for RNA viruses with very fast evolutionary rates. Target capture sequencing, as the name suggests, is a method enriching target genomes in the NGS library using complementary probes to increase the sensitivity of NGS analysis (3–5).

Typically, multiple 80- or 120-mer biotinylated DNA or RNA probes are used, which are designed to overlap so that the entire target genome is covered. The designed probes hybridize to specific target genomes, which is expected to reduce the sequence reads derived from non-target genomes such as bacterial and host-derived genomes, allowing highly sensitive detection of the target genomes. Another advantage of target capture sequencing is its high tolerance for mismatches between the probe and the target nucleotide region (9). This advantage has been useful in the search for unidentified pathogens and/or RNA viruses that frequently mutate, such as SARS-CoV-2 and influenza A viruses. To date, multiple studies have applied target capture sequencing for the genome analysis of SARS-CoV-2 (9–16). More recently, a group attempted to sequence the monkey pox virus genome using target capture sequencing (17).

Target capture sequencing also enables the detection of multiple pathogens by simultaneously incorporating a large number of probes derived from various pathogen genomes into a single assay. Target capture sequencing for viral pathogens was first applied to specific viral genome analyses (18–20), and then commercial multi-pathogen DNA or RNA probe panels, or proprietary oligonucleotide panels customizable for target pathogens, were introduced and evaluated using clinical and/or non-clinical specimens (5, 9, 11, 12, 14, 21–25). For example, ViroCap, a custom probe panel designed from genome sequences of 34 families of vertebrate DNA or RNA viruses, dramatically increased the number of sequence reads derived from the viruses, as well as their breadth of coverage and average coverage, and detected a total of 32 viruses from the clinical specimens of 22 patients (23). Thus target capture sequencing has been proven useful for both research and clinical diagnostics. However, as the efficiency of enrichment varies depending on the number of probes and targets included in the probe panel and the type of clinical specimen, as well as the library preparation method, validation is essential before application to clinical diagnostics. Moreover, it is critical to determine the limitations or detection sensitivity of each probe panel in clinical specimens with low viral load.

In this study, we first investigate the enrichment efficiency of target capture sequencing compared to metagenomic sequencing using clinical respiratory specimens collected from patients diagnosed with SARS-CoV-2 or type A influenza with sufficient copy numbers of viral genomes. We then evaluate the utility of target capture sequencing in clinical respiratory specimens, with a focus on clinical specimens containing low copy numbers of viral genomes. As a representative, commercially available comprehensive panel, we use the Twist Comprehensive Viral Research Panel (Twist Bioscience, San Francisco, CA), which contains reference sequences for a total of 15,488 different strains of 3,153 viruses, including zoonotic and human epizootic pathogens (https://www.twistbioscience.com/products/ngs/fixed-panels/comprehensive-viral-research-panel?tab=overview). Our results will provide important insights into the clinical diagnostic applications of target capture sequencing for various viral pathogens.

## Results

### Comparison of metagenomic and target capture sequencing

To evaluate the enrichment efficiency of target capture sequencing, we first selected two clinical specimens with Ct values below 30 for either SARS-CoV-2 or influenza A virus by real-time PCR: The copy number of SARS-CoV-2 in the RNA solution extracted from CS2022-0121 was 59.3 copies/µL and that of influenza A virus in the F16-31-UTM RNA solution was 544 copies/µL (Table 2).

**Table 1.**
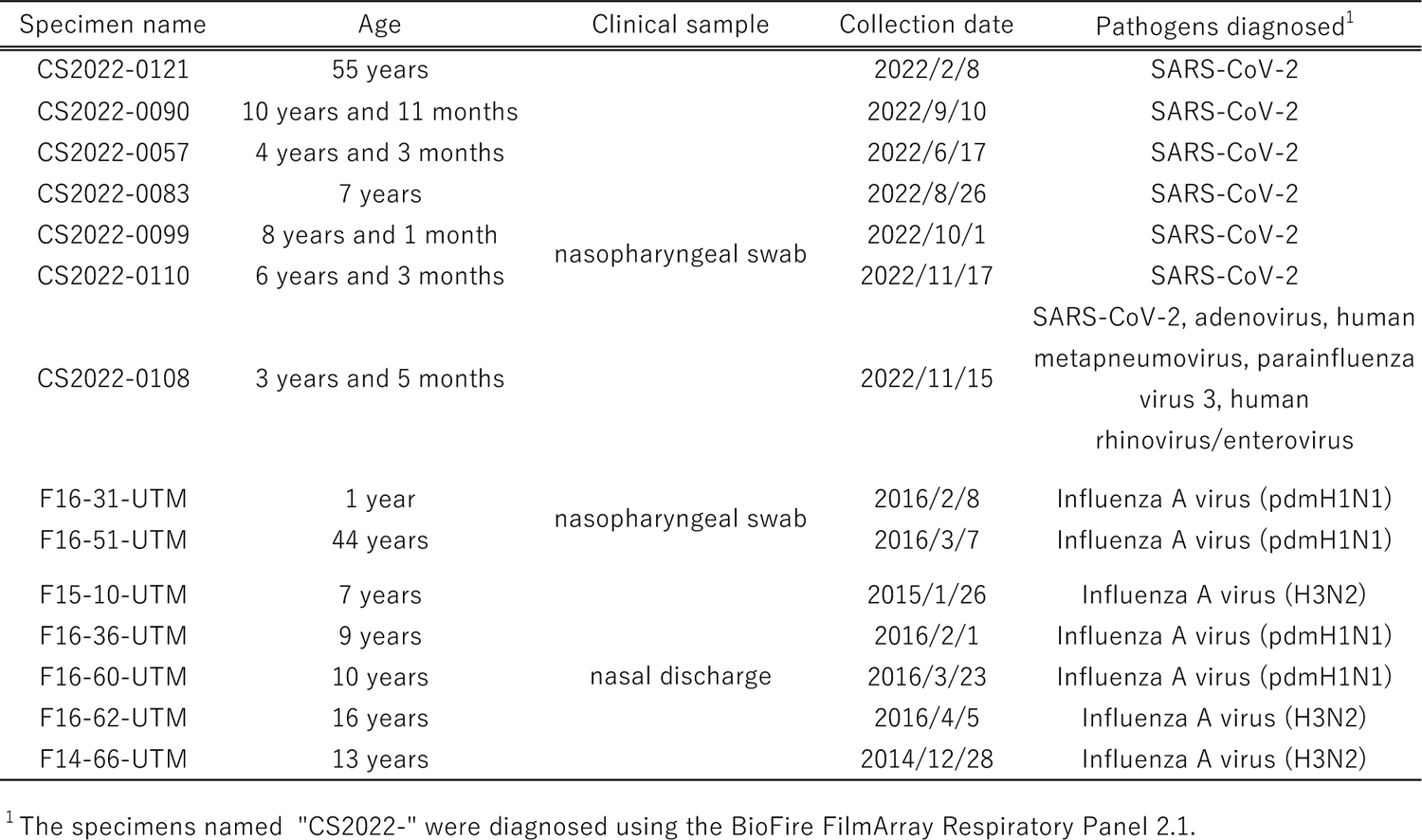
List of clinical specimens collected from patients showing respiratory diseases used in this study.

**Table 2.** Comparison of metagenomic and target capture sequencing using specimens containing SARS-CoV-2 and influenza A virus per million reads^1^.

| Specimen name | Cp value | Conc. cp/ μL | Assay | Segment | Number of mapped reads to the human genome (%) | | Mapped reads to the reference viral genome<sup>2</sup> (%) | | Average coverage | Consensus length (nt) | Breadth of coverage (%) |
| --- | --- | --- | --- | --- | --- | --- | --- | --- | --- | --- | --- |
| CS2022-0121 (SARS-CoV-2) | 28.85 | 59.33 | Metagenomic | N.A. | 988,592 | ( 98.86 ) | 40 | ( 0.00 ) | 0.2 | 5,024 | 16.8 |
|  |  |  | Metagenomic with depletion of rRNA |  | 968,871 | ( 96.89 ) | 53 | ( 0.01 ) | 0.3 | 5,160 | 17.3 |
|  |  |  | Target capture |  | 953,497 | ( 95.35 ) | 7,331 | ( 0.73 ) | 35.0 | 28,573 | 95.6 |
|  |  |  | Target capture with depletion of rRNA |  | 959,712 | ( 95.97 ) | 2,101 | ( 0.21 ) | 9.8 | 22,826 | 76.4 |
| F16-31-UTM (A(H1N1)pdm09) | 28.54 | 544 | Metagenomics | PB2 | 985,241 | ( 98.52 ) | 2 | ( 0.00 ) | 0.1 | 197 | 8.4 |
|  |  |  |  | PB1 |  |  | 0 | ( 0.00 ) | 0.0 | 0 | 0.0 |
|  |  |  |  | PA |  |  | 3 | ( 0.00 ) | 0.2 | 366 | 16.4 |
|  |  |  |  | HA |  |  | 2 | ( 0.00 ) | 0.2 | 179 | 10.1 |
|  |  |  |  | NP |  |  | 2 | ( 0.00 ) | 0.2 | 256 | 16.4 |
|  |  |  |  | NA |  |  | 0 | ( 0.00 ) | 0.0 | 0 | 0.0 |
|  |  |  |  | MP |  |  | 1 | ( 0.00 ) | 0.1 | 69 | 6.7 |
|  |  |  |  | NS |  |  | 0 | ( 0.00 ) | 0.0 | 0 | 0.0 |
|  |  |  |  | Total |  |  | 10 | ( 0.00 ) | 0.1 | 1,067 | 7.8 |
|  |  |  | Metagenomics with depletion of rRNA | PB2 | 980,868 | ( 98.09 ) | 0 | ( 0.00 ) | 0.0 | 0 | 0.0 |
|  |  |  |  | PB1 |  |  | 2 | ( 0.00 ) | 0.1 | 123 | 5.3 |
|  |  |  |  | PA |  |  | 2 | ( 0.00 ) | 0.1 | 245 | 11.0 |
|  |  |  |  | HA |  |  | 0 | ( 0.00 ) | 0.0 | 0 | 0.0 |
|  |  |  |  | NP |  |  | 2 | ( 0.00 ) | 0.2 | 300 | 19.2 |
|  |  |  |  | NA |  |  | 0 | ( 0.00 ) | 0.0 | 0 | 0.0 |
|  |  |  |  | MP |  |  | 0 | ( 0.00 ) | 0.0 | 0 | 0.0 |
|  |  |  |  | NS |  |  | 0 | ( 0.00 ) | 0.0 | 0 | 0.0 |
|  |  |  |  | Total |  |  | 6 | ( 0.00 ) | 0.1 | 668 | 4.9 |
|  |  |  | Target capture | PB2 | 956,973 | ( 95.70 ) | 4,944 | ( 0.49 ) | 306.1 | 2,323 | 99.23 |
|  |  |  |  | PB1 |  |  | 2,851 | ( 0.29 ) | 177.8 | 2,276 | 97.22 |
|  |  |  |  | PA |  |  | 3,510 | ( 0.35 ) | 228.1 | 2,185 | 97.85 |
|  |  |  |  | HA |  |  | 2,546 | ( 0.25 ) | 208.0 | 1,758 | 98.93 |
|  |  |  |  | NP |  |  | 2,332 | ( 0.23 ) | 216.6 | 1,553 | 99.23 |
|  |  |  |  | NA |  |  | 1,547 | ( 0.15 ) | 154.3 | 1,444 | 99.04 |
|  |  |  |  | MP |  |  | 1,312 | ( 0.13 ) | 186.4 | 972 | 94.64 |
|  |  |  |  | NS |  |  | 417 | ( 0.04 ) | 66.5 | 876 | 98.43 |
|  |  |  |  | Total |  |  | 19,459 | ( 1.95 ) | 207.3 | 13,387 | 98.2 |
|  |  |  | Target capture with depletion of rRNA | PB2 | 963,240 | ( 96.32 ) | 2,166 | ( 0.22 ) | 134.7 | 2,029 | 86.67 |
|  |  |  |  | PB1 |  |  | 3,105 | ( 0.31 ) | 203.5 | 2,162 | 96.82 |
|  |  |  |  | PA |  |  | 283 | ( 0.03 ) | 44.0 | 866 | 97.30 |
|  |  |  |  | HA |  |  | 1,212 | ( 0.12 ) | 120.8 | 1,434 | 98.35 |
|  |  |  |  | NP |  |  | 831 | ( 0.08 ) | 118.3 | 974 | 94.84 |
|  |  |  |  | NA |  |  | 1,379 | ( 0.14 ) | 113.1 | 1,753 | 98.65 |
|  |  |  |  | MP |  |  | 2,939 | ( 0.29 ) | 184.0 | 2,261 | 96.58 |
|  |  |  |  | NS |  |  | 727 | ( 0.07 ) | 66.8 | 1,529 | 97.70 |
|  |  |  |  | Total |  |  | 12,642 | ( 1.26 ) | 135.2 | 13,008 | 95.4 |

<sup>1</sup> All data were standardized and calculated per million reads to compare differences between assays.<sup>2</sup> hCoV-19\_Wuhan\_WIV04\_2019 (EPI\_ISL\_402124) and A/California/04/2009 (H1N1) (EPI\_ISL\_376192) were used as the reference sequences of CS2022-0121 and F16-31-UTM, respectively.

In the CS2022-0121 library, metagenomic sequencing revealed 40 viral reads mapping to SARS-CoV-2 and only 53 viral reads even after the human rRNA-removal treatment. Their consensus lengths were 5,024 and 5,160 nt covering approximately 17% of the SARS-CoV-2 genome, and their average coverages were 0.2 and 0.3, respectively. On the other hand, target capture sequencing showed dramatic improvements in all the metrics. The numbers of reads mapped to SARS-CoV-2 in target capture sequencing without and with rRNA-removal treatment were 7,331 and 2,101, respectively. These reads resulted in consensus sequences of 28,573 and 22,826 nt covering 95.6% and 76.4% of the SARS-CoV-2 genome, respectively. The enrichment efficiency of SARS-CoV-2 in the target capture sequencing assay without rRNA-removal treatment against metagenomic sequencing was approximately 183.

In the F16-31-UTM library, the data obtained in each assay were analyzed in each of the eight segments of the influenza A virus genome (the PB2, PB1, PA, HA, NP, NA, M, NS gene segments) (Table 2). Metagenomic sequencing without rRNA-removal treatment and that with rRNA-removal treatment revealed only 10 and 6 reads mapped to the influenza A virus genome, covering 7.8 and 4.9 % of the total genomes, respectively. No reads derived from the NA gene were obtained in either assay, suggesting that subtype identification was not possible with metagenomic sequencing. The number of reads mapped to the influenza A virus by target capture sequencing without rRNA-removal treatment and that with rRNA-removal treatment were 19,459 and 12,642, respectively.

These reads resulted in consensus sequences of 13,387 and 13,008 nt covering 98.2% and 95.4% of the influenza A virus genome, respectively. In both assays, the consensus lengths covering >94% were obtained for all gene segments except for the PB2 gene in the target capture sequencing with rRNA-removal treatment. The enrichment efficiency of influenza A virus in target capture sequencing assay without rRNA-removal treatment compared to metagenomic sequencing without rRNA-removal treatment was approximately 1,950.

rRNA-removal treatment had a more limited effect in the target capture sequencing compared to the metagenomic sequencing (Table 2). In metagenomic sequencing in the CS2022-0121 specimen, the percentage of reads derived from the human genome declined from 98.9% without rRNA-removal treatment to 96.9% with rRNA-removal treatment. A slight decline was observed in the F16-31-UTM specimen, from 98.5% to 98.1%. However, target capture sequencing using the same specimens did not yield a reduction in the percentage of human genomes; the percentages were approximately 95%–96% for CS2022-0121 and F16-31-UTM irrespective of rRNA removal.

Accordingly, the non-rRNA removal treat assay was adopted for all subsequent target capture sequencing. All sets of reads aligned to either the SARS-CoV-2 or influenza A virus reference genome, downsampled to 1,000,000, are shown in Supplementary Figures (Fig. S1–2).

### Availability of target capture sequencing in clinical specimens with low viral loads

To investigate the availability of target capture sequencing in clinical specimens with low viral loads, we selected clinical specimens with Ct values >30 against either SARS-CoV-2 or influenza A viruses. The SARS-CoV-2-positive specimens (CS2022-0099, −0110, −0108, −0083, −0090, and −0057) contained 0.52–9.79 RNA copies/µL of SARS-CoV-2 genome (Table 3), and the influenza A virus (A(H1N1)pdm09 or H3N2 subtypes)-positive specimens (F16-51-UTM, F16-36-UTM, F16-60-UTM, F15-10-UTM, F16-62-UTM, and F14-66-UTM) contained 1.63–25 RNA copies/µL of influenza A virus genome (Table 4).

**Table 3.** Overview of the genome reads per million reads obtained from the clinical samples showing high Ct values against SARS-CoV-2 (> 30) using target capture sequencing.

| Specimen name | Ct values | Conc. cp/$\mu$ L | Mapped reads to reference viral genome<sup>1</sup> | | Average coverage | Consensus length (nt) | Breadth of coverage (%) |
| --- | --- | --- | --- | --- | --- | --- | --- |
| CS2022-0099 | 31.66 | 9.79 | 2,562 | ( 0.26 ) | 12.49 | 23904 | 80.0 |
| CS2022-0110 | 32.14 | 6.88 | 7,845 | ( 0.78 ) | 26 | 23980 | 80.2 |
| CS2022-0108 | 33.95 | 0.52 | 7 | ( 0.00 ) | 0.02 | 395 | 1.3 |
| CS2022-0083 | 34.52 | 1.55 | 1,826 | ( 0.18 ) | 8.23 | 4704 | 15.7 |
| CS2022-0090 | 34.59 | 0.86 | 6 | ( 0.00 ) | 0.02 | 198 | 0.7 |
| CS2022-0057 | 34.65 | 0.52 | 14 | ( 0.00 ) | 0.04 | 305 | 1.0 |

<sup>1</sup> hCoV-19\_Wuhan\_WIV04\_2019 (EPI\_ISL\_402124) was used as a reference sequence.

**Table 4.**
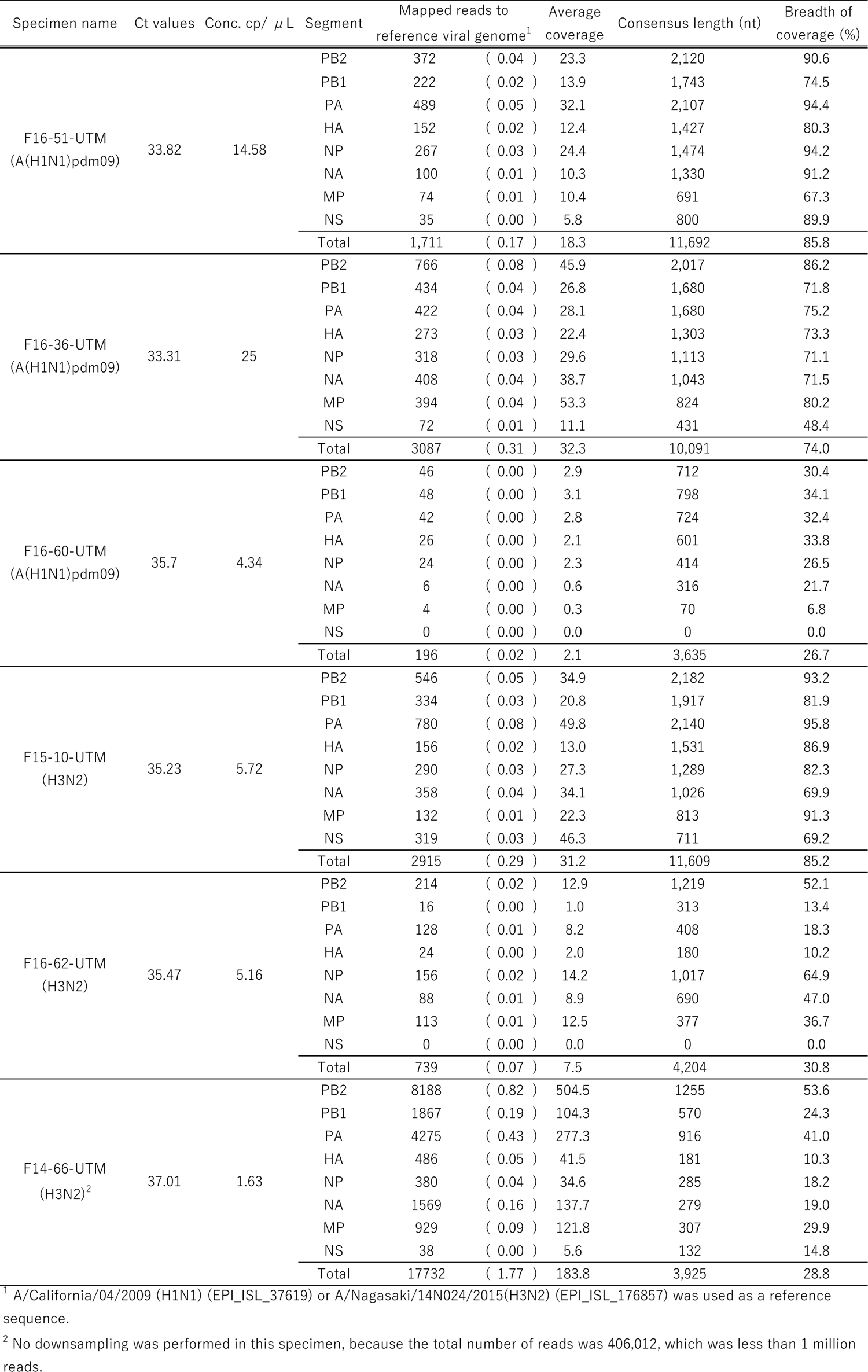
Overview of the genome reads per million reads obtained from the clinical specimens showing high Ct values against Influenza A virus (> 30)

| Specimen name | Ct values | Conc. cp/ $\mu$ L | Segment | Mapped reads to<br>reference viral genome<sup>1</sup> | | Average<br>coverage | Consensus length (nt) | Breadth of<br>coverage (%) |
| --- | --- | --- | --- | --- | --- | --- | --- | --- |
| F16-51-UTM<br>(A(H1N1)pdm09) | 33.82 | 14.58 | PB2 | 372 | ( 0.04 ) | 23.3 | 2,120 | 90.6 |
|  |  |  | PB1 | 222 | ( 0.02 ) | 13.9 | 1,743 | 74.5 |
|  |  |  | PA | 489 | ( 0.05 ) | 32.1 | 2,107 | 94.4 |
|  |  |  | HA | 152 | ( 0.02 ) | 12.4 | 1,427 | 80.3 |
|  |  |  | NP | 267 | ( 0.03 ) | 24.4 | 1,474 | 94.2 |
|  |  |  | NA | 100 | ( 0.01 ) | 10.3 | 1,330 | 91.2 |
|  |  |  | MP | 74 | ( 0.01 ) | 10.4 | 691 | 67.3 |
|  |  |  | NS | 35 | ( 0.00 ) | 5.8 | 800 | 89.9 |
|  |  |  | Total | 1,711 | ( 0.17 ) | 18.3 | 11,692 | 85.8 |
| F16-36-UTM<br>(A(H1N1)pdm09) | 33.31 | 25 | PB2 | 766 | ( 0.08 ) | 45.9 | 2,017 | 86.2 |
|  |  |  | PB1 | 434 | ( 0.04 ) | 26.8 | 1,680 | 71.8 |
|  |  |  | PA | 422 | ( 0.04 ) | 28.1 | 1,680 | 75.2 |
|  |  |  | HA | 273 | ( 0.03 ) | 22.4 | 1,303 | 73.3 |
|  |  |  | NP | 318 | ( 0.03 ) | 29.6 | 1,113 | 71.1 |
|  |  |  | NA | 408 | ( 0.04 ) | 38.7 | 1,043 | 71.5 |
|  |  |  | MP | 394 | ( 0.04 ) | 53.3 | 824 | 80.2 |
|  |  |  | NS | 72 | ( 0.01 ) | 11.1 | 431 | 48.4 |
|  |  |  | Total | 3087 | ( 0.31 ) | 32.3 | 10,091 | 74.0 |
| F16-60-UTM<br>(A(H1N1)pdm09) | 35.7 | 4.34 | PB2 | 46 | ( 0.00 ) | 2.9 | 712 | 30.4 |
|  |  |  | PB1 | 48 | ( 0.00 ) | 3.1 | 798 | 34.1 |
|  |  |  | PA | 42 | ( 0.00 ) | 2.8 | 724 | 32.4 |
|  |  |  | HA | 26 | ( 0.00 ) | 2.1 | 601 | 33.8 |
|  |  |  | NP | 24 | ( 0.00 ) | 2.3 | 414 | 26.5 |
|  |  |  | NA | 6 | ( 0.00 ) | 0.6 | 316 | 21.7 |
|  |  |  | MP | 4 | ( 0.00 ) | 0.3 | 70 | 6.8 |
|  |  |  | NS | 0 | ( 0.00 ) | 0.0 | 0 | 0.0 |
|  |  |  | Total | 196 | ( 0.02 ) | 2.1 | 3,635 | 26.7 |
| F15-10-UTM<br>(H3N2) | 35.23 | 5.72 | PB2 | 546 | ( 0.05 ) | 34.9 | 2,182 | 93.2 |
|  |  |  | PB1 | 334 | ( 0.03 ) | 20.8 | 1,917 | 81.9 |
|  |  |  | PA | 780 | ( 0.08 ) | 49.8 | 2,140 | 95.8 |
|  |  |  | HA | 156 | ( 0.02 ) | 13.0 | 1,531 | 86.9 |
|  |  |  | NP | 290 | ( 0.03 ) | 27.3 | 1,289 | 82.3 |
|  |  |  | NA | 358 | ( 0.04 ) | 34.1 | 1,026 | 69.9 |
|  |  |  | MP | 132 | ( 0.01 ) | 22.3 | 813 | 91.3 |
|  |  |  | NS | 319 | ( 0.03 ) | 46.3 | 711 | 69.2 |
|  |  |  | Total | 2915 | ( 0.29 ) | 31.2 | 11,609 | 85.2 |
| F16-62-UTM<br>(H3N2) | 35.47 | 5.16 | PB2 | 214 | ( 0.02 ) | 12.9 | 1,219 | 52.1 |
|  |  |  | PB1 | 16 | ( 0.00 ) | 1.0 | 313 | 13.4 |
|  |  |  | PA | 128 | ( 0.01 ) | 8.2 | 408 | 18.3 |
|  |  |  | HA | 24 | ( 0.00 ) | 2.0 | 180 | 10.2 |
|  |  |  | NP | 156 | ( 0.02 ) | 14.2 | 1,017 | 64.9 |
|  |  |  | NA | 88 | ( 0.01 ) | 8.9 | 690 | 47.0 |
|  |  |  | MP | 113 | ( 0.01 ) | 12.5 | 377 | 36.7 |
|  |  |  | NS | 0 | ( 0.00 ) | 0.0 | 0 | 0.0 |
|  |  |  | Total | 739 | ( 0.07 ) | 7.5 | 4,204 | 30.8 |
| F14-66-UTM<br>(H3N2)<sup>2</sup> | 37.01 | 1.63 | PB2 | 8188 | ( 0.82 ) | 504.5 | 1255 | 53.6 |
|  |  |  | PB1 | 1867 | ( 0.19 ) | 104.3 | 570 | 24.3 |
|  |  |  | PA | 4275 | ( 0.43 ) | 277.3 | 916 | 41.0 |
|  |  |  | HA | 486 | ( 0.05 ) | 41.5 | 181 | 10.3 |
|  |  |  | NP | 380 | ( 0.04 ) | 34.6 | 285 | 18.2 |
|  |  |  | NA | 1569 | ( 0.16 ) | 137.7 | 279 | 19.0 |
|  |  |  | MP | 929 | ( 0.09 ) | 121.8 | 307 | 29.9 |
|  |  |  | NS | 38 | ( 0.00 ) | 5.6 | 132 | 14.8 |
|  |  |  | Total | 17732 | ( 1.77 ) | 183.8 | 3,925 | 28.8 |

<sup>1</sup> A/California/04/2009 (H1N1) (EPI\_ISL\_37619) or A/Nagasaki/14N024/2015(H3N2) (EPI\_ISL\_176857) was used as a reference sequence.
<sup>2</sup> No downsampling was performed in this specimen, because the total number of reads was 406,012, which was less than 1 million reads.

For SARS-CoV-2-positive specimens, a slight positive correlation was observed between RNA copy number per microliter and breadth of coverage (R²=0.48) (Fig. 1A, Table 3). Target capture sequencing identified viral reads covering approximately 80% of the SARS-CoV-2 genome with an average coverage of >12 in CS2022-0099 and CS2022-0110, which contained >6 RNA copies/µL. In contrast, the specimens containing <5 RNA copies/µL, with the exception of CS2022-0083, showed only 1–14 viral reads derived from SARS-CoV-2 covering 0.7% to 1.3% of the SARS-CoV-2 genome. Interestingly, CS2022-0083 showed 1,826 viral reads covering 15.7% of the total length, despite containing only 1.55 RNA copies/µL of SARS-CoV-2.

**Figure 1.**
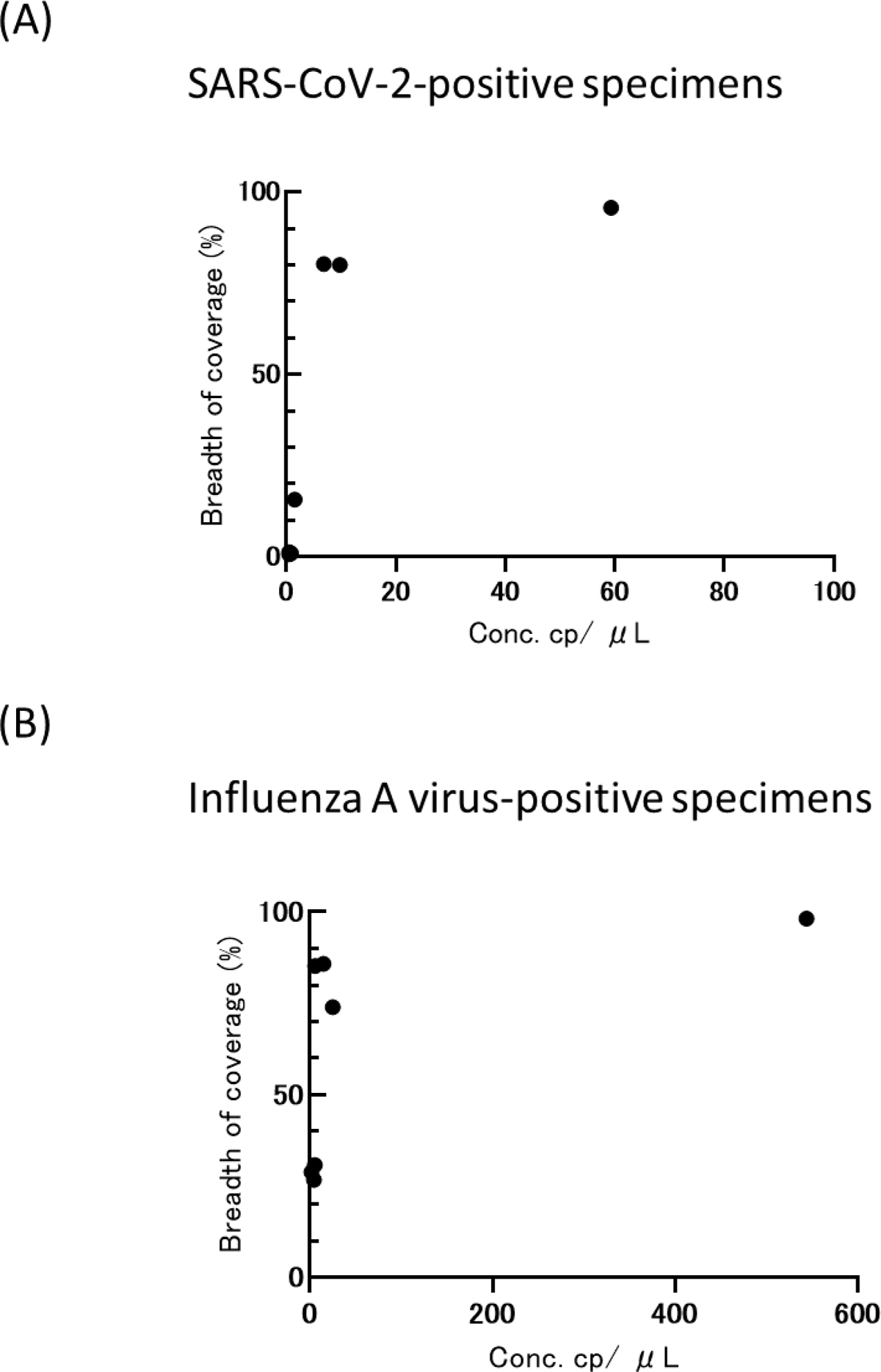
Correlation between RNA copy number per microliter and breadth of coverage (%) obtained by using target capture sequencing in RNA extracts from clinical specimens containing SARS-CoV-2 (A) and influenza A virus genomes (B), respectively.

For influenza A virus-positive specimens, there was no clear positive correlation between RNA copy number per microliter and breadth of coverage (R²=0.29) (Fig 1B, Table 4). In specimens, F16-51-UTM and F16-36-UTM, which contained >14 RNA copies of the influenza A virus, viral reads covering >70% of the influenza A virus genome were obtained with an average coverage >18. Except for the NS gene in F16-36-UTM, >67% breadth of coverage was obtained in all segments in these specimens, suggesting efficient genomic analysis by targeted capture sequencing. Despite containing only 5.72 RNA copies of influenza A virus, the F15-10-UTM specimen yielded a total of 2,915 viral reads covering 85.2% of the influenza A virus genome. Surprisingly, in specimens containing less than ∼5 RNA copies/uL, i.e., F16-60-UTM, F16-62-UTM, and F4-66-UTM, consensus sequences covering about 26%–30% of the influenza A virus genome were obtained. The breadth of coverage obtained in HA and NA genes of these specimens exceeded 10%, which was sufficient for identification of their viral subtypes. In these analyses, although one-step real-time PCR was used to quantify RNA copies of influenza A virus, it was confirmed that there was a correlation between real-time PCR and digital PCR quantification values in some specimens (data not shown).

### Detection of simultaneous infections with multiple viral pathogens

Viral pathogen genomes other than SARS-CoV-2 were detected in two of the SARS-CoV-2-positive specimens, CS2022-0083 and CS2022-0108. No viral pathogen genomes other than influenza A virus were detected in influenza A virus-positive specimens. In CS2022-0083, 1,553 of the obtained reads were mapped to Circovirus-like genome DCCV-4 (accession number NC_030470.1) with 53.2% breadth of coverage and 63.4 average coverage (Table 5). Unfortunately, for the circovirus-like genome DCCV-4, only two genome sequences obtained directly from environmental samples from a freshwater lake in China have been published, so it was not possible to estimate whether the viruses with similar genomes affected respiratory symptoms in this study. In the CS2022-0108 specimen, which was positive for adenovirus (ADV), human metapneumovirus (MNV), parainfluenza virus 3 (PIV3), and human rhinovirus(RV)/enterovirus(EV) in addition to SARS-CoV-2 by diagnosis with the BioFire FilmArray Respiratory Panel 2.1 (Table 1), 4,354 of the obtained reads were mapped to the reference MN173594.1, which was the enterovirus D68 strain USA/2018/CA-RGDS-1056 polyprotein gene with 93.2% breadth of coverage and 88.5 average coverage (Table 5). BLAST analysis revealed that a consensus sequence of 7,335 nt had the highest identity (99.4%) with JH-EV-50/2022 (accession number OP572066.1) isolated in the USA in 2022. Although it was below the threshold for taxonomy analysis, 30 reads mapped to PIV3 (accession number NC_038270.1) and 28 reads mapped to ADV (accession number NC_001405.1) were found, respectively. However, no reads mapped to MNV, so the copy number of MNV may have been below the detection limit of the target capture sequencing used in this study.

**Table 5.** Detection of simultaneous viral infections by the Find Best References tool with the Clustered Reference Viral DataBase^1^.

| Specimen name | Input reads | Number of reads mapped | Breadth of coverage (%) | Average coverage | Accession no. of best match reference | Reference length (nt) | Taxonomy (definition) |
| --- | --- | --- | --- | --- | --- | --- | --- |
| CS2022-0083 | 1,905,260 | 1,553 | 53.2 | 63.4 | NC_030470.1 | 2,985 | Circovirus-like genome DCCV-4, complete genome |
| CS2022-0108<sup>2</sup> | 2,016,746 | 4,354 | 93.2 | 88.5 | MN173594.1 | 7,272 | Enterovirus D68 strain USA/2018/CA-RGDS-1056 polyprotein gene, complete cds. |

<sup>1</sup> The Clustered Reference Viral DataBase (RVDB, v21.0 June 2021) was used as the reference in the Genomics Workbench software. The thresholds of minimum number of reads on a reference, minimum coverage, and minimum fraction of reference covered were set to be 100, 10, and 0.5, respectively.
<sup>2</sup> The BioFire FilmArray Respiratory Panel 2.1 detected adenovirus, human metapneumovirus, parainfluenza virus 3, and human rhinovirus/enterovirus in addition to SARS-CoV-2 in this specimen.

## Discussion

Here, we have demonstrated the effectiveness of target capture sequencing directly from respiratory clinical specimens with low viral load. The capture panel, the Twist Comprehensive Viral Research Panel, was selected for its ability to capture a very wide range of viral pathogens; to our knowledge, this is the first evaluation using human respiratory clinical specimens with low viral load although the panel has been used on human cerebrospinal fluid (26), saliva, blood and feces from wild bats (27) and mosquito samples (28). Target capture sequencing with this panel yielded approximately 180- and 2000-fold higher read counts of SARS-CoV-2 or influenza A virus, respectively, than metagenomic sequencing when RNA extracts from specimens containing 59.3 or 544 RNA copies/µL of SARS-CoV-2 or influenza A virus were used, respectively. Although library preparation conditions, sequencers and capture panels were different, previous studies also found that the target capture sequencing had higher sensitivity than metagenomic sequencing (3, 4, 9, 11–13, 16, 22, 23, 29, 30), reinforcing our present results.

The detection sensitivity of the target capture sequencing may not differ substantially from that of RT-PCR methods, although a variety of experimental conditions and positivity thresholds, such as NGS metrics (breadth of coverage, average coverage, etc.), have been adopted in the target capture sequencing studies (15, 16, 29). In fact, all specimens positive for SARS-CoV-2 or influenza type A yielded the corresponding virus-derived reads in our target capture sequencing, although SARS-CoV-2-positive specimens with Ct values >35 were not available in this study. Nagy-Szakal et al. reported that the positive and negative percentage concordances between the RT-PCR method and target capture sequencing using nasopharyngeal swab specimens were 96.7% and 100%, respectively (15). On the other hand, the enrichment efficiency was clearly higher for influenza A virus-positive specimens than for SARS-CoV-2-positive specimens. The Twist Comprehensive Viral Research Panel contains probes derived from 8,050 strains of influenza A viruses and only one strain of SARS-CoV-2 (strain name unavailable). Therefore, there may be a greater number of probes hybridizing to the influenza A virus genome than to the SARS-CoV-2 genome. Unfortunately, the detailed design of the panel, which could be crucial for enrichment (11, 21, 31), was not available, and thus it was not clear how the panel affected our NGS results.

There was no clear correlation between the RNA copy number of the viral genome and the number of corresponding reads obtained from the clinical specimens with low viral load. Interestingly, F15-10-UTM, which contained only 5.72 copies/μL of the influenza A virus genome, yielded viral reads that covered 85% of the H3N2 influenza virus, while F16-60-UTM and F16-62-UTM, which had similar copy numbers, yielded only 26.7% and 30.8% breadths of coverage, respectively.

Likewise, it has been reported that the clinical specimens with Ct values >30 showed a variety of enrichment efficiencies after target capture sequencing even among the specimens with the same Ct values (11, 12, 15). On the other hand, in NGS libraries prepared from the dilution of cultured viruses rather than clinical specimens (13), there seems to be a clearer correlation between the number of viral reads and genome copies. Thus, it may be difficult to obtain a stable number of reads from NGS libraries prepared from clinical specimens with low viral load, as the methods of specimen collection and the proportion of host- or bacteria-derived genomes are quite different in each specimen.

As shown in our study and in previous studies (16, 23, 29), the target capture sequencing using probes for multiple pathogens is also very useful for the identification of co-infections. Almost the whole genome of EV-D68 was obtained from the CS2022-0108 specimen. EV-D68 is an emerging viral pathogen first identified in 1962 in hospitalized children with respiratory disease (32). Since 2005, several countries have reported an increase in the number of patients with respiratory diseases caused by EV-D68 (33). In addition, a rapid increase in EV-D68 infections was reported from eight European countries in 2021 (34). In Japan, EV-D68 outbreaks have been reported several times (35–37). Interestingly, EV-D68 infections were reported in 25 (14%) of 197 specimens positive for HRV/EV by BIOFIRE® Respiratory 2.1 collected from 1 September to 13 October 2022 at a hospital in Tokyo, Japan (different from the hospital in this study) (IASR Vol. 43 pp. 290–291: 2022, Dec. https://www.niid.go.jp/niid/ja/diseases/a/ev-d68/2335-idsc/iasr-news/11650-514p01.html) (in Japanese). These hospitals are located close to each other in Tokyo and the specimens were collected during the same season, suggesting that an epidemic of EV-D68 occurred in Tokyo along with the SARS-CoV-2 epidemic. These results suggest that target capture sequencing with comprehensive probe panels could not only detect a wide range of causative viruses, but also contribute to epidemiological studies.

Greater probe diversity allows the detection of many targeted genomes in target capture sequencing, but there may be a trade-off in increasing the number of off-target reads (11). Target capture sequencing dramatically increased the number of target viral reads, but human genome reads still accounted for >95% of the total reads in this study. In addition, unfortunately, human rRNA-removal treatment did not significantly reduce the percentage of human genome reads, but instead reduced the number of target viral reads in target captured libraries. Other pre-treatments, such as filtering of original specimens and post-extraction DNase treatment, also showed little effect on target capture efficiency (29). Therefore, in our study, especially for specimens with low viral load, we decided to use only high-speed centrifugation prior to RNA extraction. However, the large number of host genomes included in the library after hybridization suggests the possibility of further improvements to increase enrichment efficiency, even when using a comprehensive probe panel.

Target capture sequencing with the comprehensive panel could be a useful tool to simultaneously identify a variety of viruses in one assay. Although recently the genetic detection of pathogens has often been done by PCR, the number of assays must be increased according to the increase in the number of targets. This increases the burden of the assay, number of processes, and management of reagents, etc. In addition, PCR methods may miss concurrent infections by other pathogens, as no further diagnosis is done if positivity for a particular pathogen is detected. Another advantage of target capture sequencing with a comprehensive panel is that it can even identify viruses that are closely related to known pathogens, thus contributing to the discovery of new viruses with pandemic potential. Our results show that the potential for identifying small amounts of pathogens is very high, although care should be taken to avoid false positives due to contamination. It is anticipated that this new technology will be further developed and applied to pathogen diagnosis and rapid response to emerging infectious disease threats.

## Materials and Methods

### Sample preparation and RNA extraction

We used 14 clinical specimens (nasopharyngeal swab or nasal discharge) collected from patients with respiratory symptoms at Showa General Hospital, Tokyo, to evaluate the target capture sequencing (Table 1). Among the 14 specimens, 7 specimens were designated CS2022- and found to be positive for SARS-CoV-2 using a BIOFIRE® Respiratory 2.1 panel (BioFire Diagnostics, Salt Lake City, UT). The remaining 7 specimens tested positive for type A influenza by real-time RT-PCR at the National Institute of Infectious Diseases, Japan, as described below. All specimens were placed in sterile tubes containing viral transport medium and stored at −80℃ until use. Prior to RNA extraction, all clinical specimens were centrifuged at 16,000g for 2 min to reduce the risk of contamination from host and bacterial-derived materials. Sixty microliters of RNA was extracted from 140 μl of each clinical specimen by using a Viral RNA Mini kit (Qiagen, Hilden, Germany) with the automated extraction platform QIAcube (Qiagen).

Real-time RT-PCR and digital PCR for identification and quantification of SARS-CoV-2 and influenza A virus Identification of SARS-CoV-2 or influenza A virus was performed by a one-step real-time RT-PCR as previously described (38, 39). The absolute number of SARS-CoV-2 RNA copies present in each extracted RNA was determined by a digital PCR using Absolute Q™ 1-step RT-dPCR Master Mix (Thermo Fisher Scientific, Waltham, MA) in a QuantStudio Absolute Q digital PCR system (Thermo Fisher Scientific). Nine microliters of a reaction mixture containing 2.97 µL of each extracted RNA was loaded. Thermal cycling was performed as follows: reverse transcription at 55℃ for 10 min, preheating at 96℃ for 10 min, and 40 cycles of denaturation at 96℃ for 5 sec and annealing/extension at 60℃ for 30 sec. The absolute number of influenza A virus RNA copies present in each extracted RNA was determined by a one-step real-time PCR based on the number of the Twist Synthetic Influenza H3N2 RNA control (Twist Bioscience). Primers and probes targeting N genes of SARS-CoV-2 (N2 assay) (38) or M genes of influenza A virus (39) were used in each PCR assay.

### Library preparation

Libraries for metagenomic sequencing of CS2022-0121 and F16-31-UTM were prepared using a NEBNext Ultra II RNA Library Prep Kit for Illumina (NEB, Ipswich, MA) to elucidate the enrichment efficiency of the target capture sequencing. Briefly, 13 uL of each RNA was converted to single-stranded cDNA using random primers after heat fragmentation, and then double-stranded cDNA was synthesized. After end-repairing and dA-tailing reactions, the adaptors diluted at a 1:10 ratio were ligated according to the NEB protocol. After size selection performed using AMPure XP beads (Beckman Coulter, Indianapolis, IN), the adaptor-ligated DNA was amplified by 17 cycles of PCR.

Libraries for the target capture sequencing were prepared using a Twist Comprehensive Viral Research Panel (Twist Bioscience) as follows. Fifteen microliters of each RNA was converted to cDNA using Protoscript II First Strand cDNA synthesis (NEB) and random primer 6 (NEB). The NEBNext Ultra II Non Directional RNA Second Strand Synthesis Module was subsequently used to convert single-stranded cDNA to double-stranded cDNA. The libraries were then generated using a Twist Library Preparation EF Kit 2.0 and Unique Dual Indices (UDI) (Twist Bioscience). Standard hybridization workflow target capture using the Twist Comprehensive Viral Research Panel was followed by a Twist Standard Target Enrichment workflow with slight modifications. Briefly, 1 μg of each dual-indexed library was dried using a vacuum concentrator with no heat. Hybridization capture was performed by adding 1 μg of the Twist Comprehensive Viral Research Panel to each library for 16 h at 70℃. After the hybridization was complete, the hybridized libraries were collected using streptavidin beads. The streptavidin binding bead slurry was then amplified according to the following protocol: initialization at 40℃ for 45 sec, followed by 21 cycles of denaturation at 98℃ for 15 sec and annealing at 60℃ for 30 sec, with a final extension at 72℃ for 30 sec. In order to validate our workflow for target capture sequencing, we preliminarily prepared the NGS libraries of the synthetic influenza H3N2 RNA control (Twist Bioscience) with two different dilutions, resulting in 10 and 1,000 RNA copies/μl, spiked into a background of human reference RNA (Agilent Technologies, Palo Alto, CA) according to the manufacturer’s instructions. After 16 h of hybridization with the Twist Comprehensive Viral Research Panel, we confirmed that sufficient amounts of sequence reads derived from influenza A virus were obtained in both samples (data not shown).

In the preparation of NGS libraries for the metagenomic and target capture sequencings of CS2022-0121 and F16-31-UTM, the effect of human rRNA removal was also assessed using a QIAseq FastSelect rRNA removal kit (Qiagen). The rRNA removal reaction was performed after RNA thermal denaturation at 95℃ 5 min according to the instructions.

### Sequencing and data analysis

The concentrations of metagenomic or target enrichened libraries were measured using Qubit 2.0 Fluorometer in combination with Qubit dsDNA HS Assay Kit (Thermo Fisher Scientific), then were analyzed on Agilent 4150 TapeStation in combination with Agilent D1000 ScreenTape System (Agilent Technologies). The library pool (up to 8 samples) which were individually diluted to 1 nM was sequenced with 150 bp Paired end reads on the Illumina MiSeq platform, using a MiSeq Reagent kit v2 (Illumina, San Diego, CA).

Generated sequence reads were imported into the Genomics Workbench software (version 21.0.4; Qiagen). The data analysis workflow was as follows. Briefly, the reads with low-quality and <30 bp reads were trimmed and downsampled to 1,000,000 reads for performing comparisons between assays or between specimens. Then, the downsampled reads were mapped to the SARS-CoV-2 (hCoV-19_Wuhan_WIV04_2019 (EPI_ISL_402124)) or influenza A viruses (A/California/04/2009 (H1N1) (EPI_ISL_376192) or A/Nagasaki/14N024/2015(H3N2) (EPI_ISL_176857) using the default parameters in the Map Reads to Reference tool. Average coverages, number of reads, and breadth of coverage against each reference sequence (>x1), and so on, were evaluated. The enrichment efficiency of target capture sequencing was calculated as the ratio of the number of reads mapped to the reference sequence by target capture sequencing without rRNA removal treatment to those by metagenomic sequencing without rRNA removal treatment.

We further performed taxonomic analysis to evaluate the target capture sequencing as a tool for detection of simultaneous infections with multiple viral pathogens. The analysis was carried out using the Find Best References using Read Mapping Tool with Clustered Reference Viral DataBase (RVDB, v21.0 June 2021) (40) as the reference in the Genomics Workbench software. The thresholds for the minimum number of reads on a reference, minimum coverage, and minimum fraction of reference covered (minimum fraction of the reference sequence to be covered by read for a reference) were set to be 100, 10, and 0.5, respectively, and the other parameters were the default settings. Then, consensus sequences were obtained by using the Map Reads to Reference tool with default settings using appropriate references.

## Ethical considerations

This study was approved by the Research and Ethical Committee of the National Institute of Infectious Diseases (NIID), Japan (approval #1544).

## Acknowledgements

This research was supported by a grant from AMED (no. JP22fk0108543) and a JSPS KAKENHI grant (no. JP22K10501). The authors declare no conflicts of interest associated with this manuscript.

## Supplemental figures

**Figure S1.**
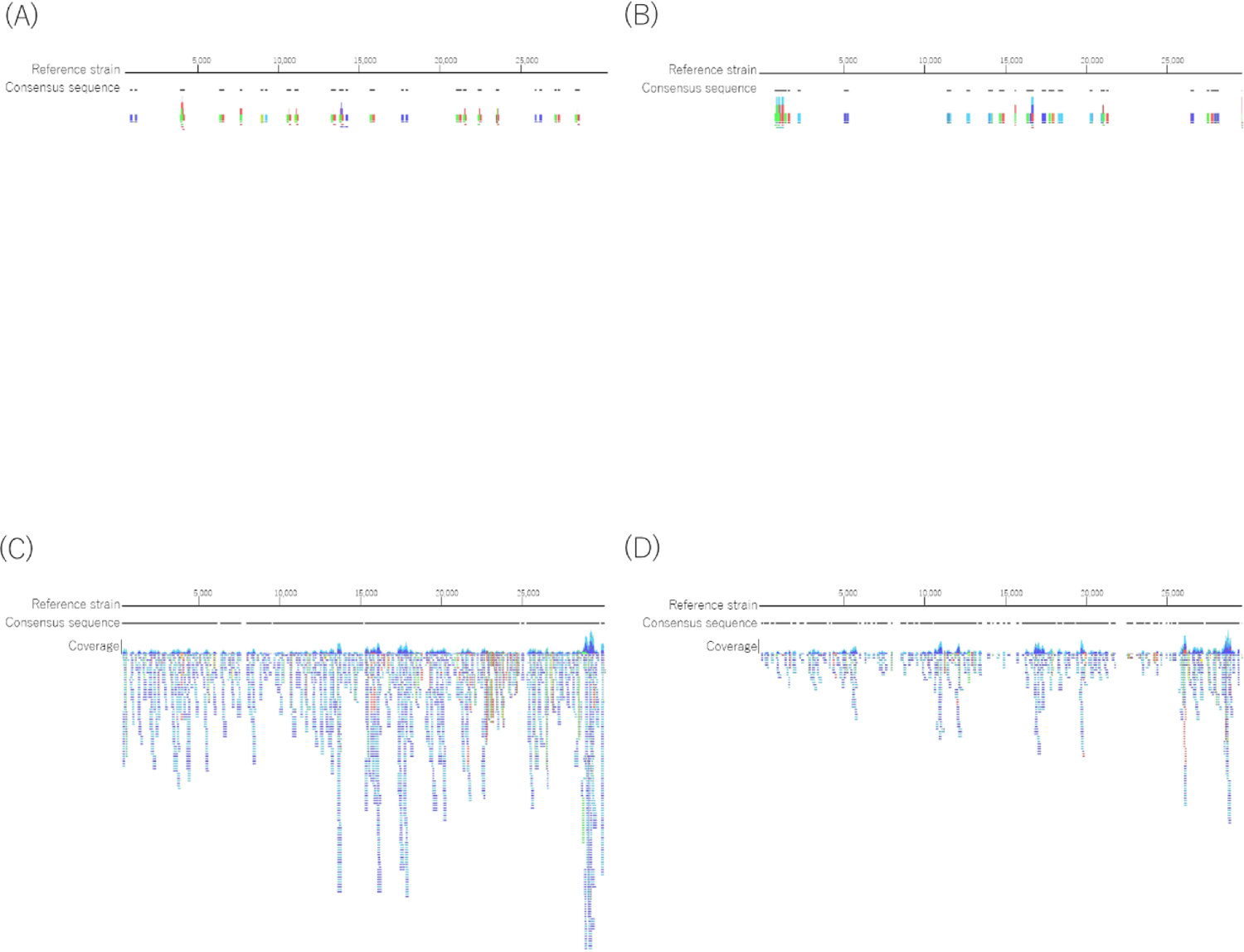
Viral reads mapped to SARS-CoV-2 obtained in the libraries of the CS2022-0121 specimen by metagenomic sequencing (A), metagenomic sequencing after depletion of human rRNA (B), target capture sequencing (C), and target capture sequencing after depletion of human rRNA are shown (D). The reference strain and consensus sequences are shown in black lines. Single reads mapping in their forward direction and reverse direction are indicated with green and red lines, respectively. The paired reads are shown with blue lines. Mismatches between the reads and reference are shown as narrow vertical traits. The coverage at each position is shown by the height of the packed reads on the read tracks. Viral reads mapped with more than 120 lines were omitted in C.

**Figure S2.**
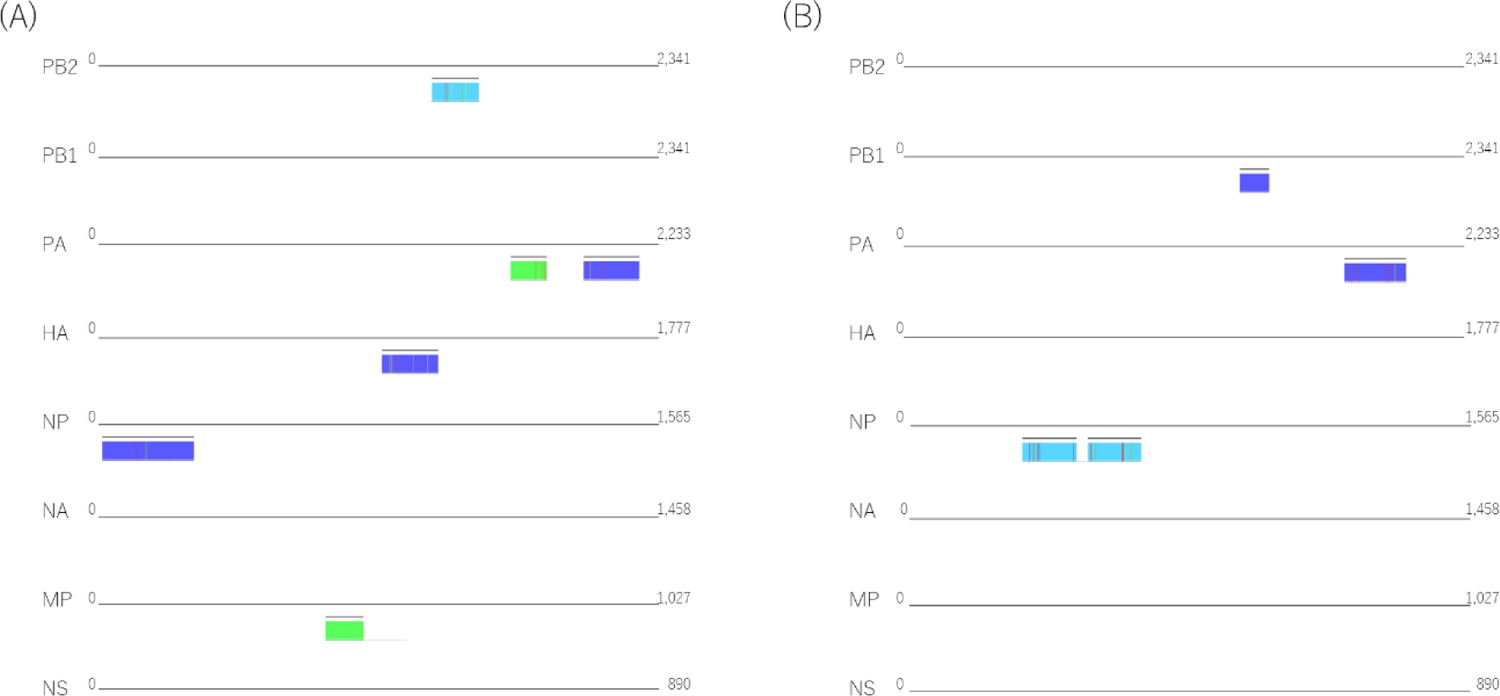
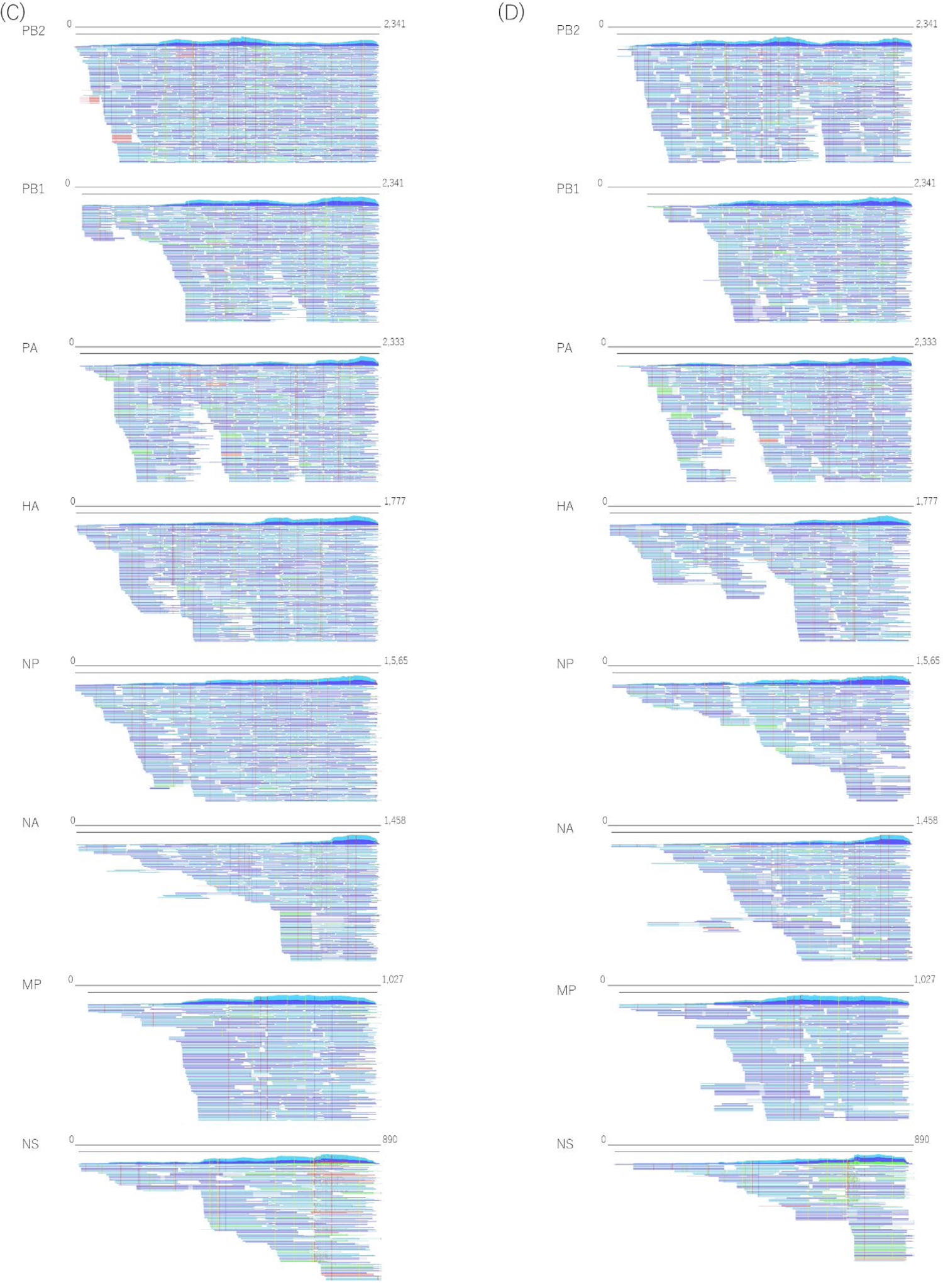
Viral reads mapped to each genome of influenza A virus obtained in the libraries of the F16-31-UTM specimen by metagenomic sequencing (A), metagenomic sequencing after depletion of human rRNA (B), target capture sequencing (C), and target capture sequencing after depletion of human (D) are shown. The reference strain and consensus sequences are indicated with black lines. Single reads mapping in their forward direction and reverse direction are shown by green and red lines, respectively. The paired reads are shown with blue lines. Mismatches between the reads and reference are shown as narrow vertical traits. The

